# Purification and partial characterization of a high hemagglutinating chitin-binding lectin from *Aponogeton natans* tubers

**DOI:** 10.1101/2021.01.21.427601

**Authors:** Shashikanth Dara, Harikrishna Naik Lavudi, Venkateswara Rao, Nanibabu Badithi, Seshagirirao Kottapalli

**Affiliations:** Department of Plant Sciences, School of Life Sciences, University of Hyderabad, Hyderabad - 500046, India; CSIR - Center for Cellular and Molecular Biology, Hubsiguda, Hyderabad – 500007, India

**Keywords:** *Aponogeton natans*, Chitin-binding lectin, hemagglutination, mitogenic proliferation

## Abstract

A novel chitin-binding lectin was isolated from the tubers of a plant *Aponogeton natans* from the monocot family Aponogetonaceae, designated as ANTL (*Aponogeton natans* tuber lectin). The lectin agglutinated both untreated and trypsin-treated rabbit erythrocytes, as well as human blood cells of groups A, B and O with different specificities. Lectin activity is inhibited by the oligomers of *N*-acetylglucosamine. ANTL is a dimeric glycoprotein with molecular weight of ∼66 KDa and has two identical sub-units of 33 KDa. The carbohydrate percent is 8.2% of the total lectin. The lectin was thermo stable up to 50°C with broad pH optima (pH 4–10). ANTL is found to be potent mitogen for normal murine and human lymphocytes at the concentration as low as 1 µg/ml. Cytotoxic studies of the lectin on human U 266 cell lines has revealed that there is 50% decrease in the proliferation. The confirmation of both the hemagglutination and mitogenic proliferation activity suggests that ANTL is a Chitin-binding lectin with diverse functions. The pharmacological relevance of ANTL as a potent mitogen with some cytotoxic effect in certain cell lines are reported for the first time.

**Highlights:**

- A novel chitin-binding lectin was purified from the tuber extracts of *Aponogeton natans* (ANTL) in a single step on chitin column by affinity chromatography.
- ANTL is a dimeric glycoprotein with a molecular weight of ∼66 KDa with high hemagglutination activity towards rabbit erythrocytes.
- The cytotoxic effect of ATNL on human cell lines U266 has shown 50 % inhibition of their proliferation.
- ANTL displayed potent mitogenic response towards murine and human peripheral blood mononuclear cells.

## 1. Introduction

Lectins are heterogenous group of glycoproteins which bind specifically and reversibly to carbohydrates or carbohydrate moieties of glycoconjugates [1]. Plant lectins are carbohydrate-binding proteins in nature they are widely distributed in seeds, roots, sap, fruits, flowers, barks, stem and leaves [2]. They play various roles in plants including packaging and transport of carbohydrates and proteins. Lectins are involved in defense mechanism against the attack of fungi, insects, pest and virus [3,4]. Lectins are used in wide range of applications such as mitogenic stimulation of lymphocytes, insulin like effect on fat cells, inhibition of fungal, bacterial, viral growth and agglutination of erythrocytes, Lectins also display insecticidal property, nuclease activity and anti-HIV property [5–7]. Most of the lectins isolated are from the dicots particularly *Leguminaceae* family. Lectins from monocots were less studied. Ever since the purification of Wheat germ agglutinin (WGA) from Gramineae, screening and isolation of lectins has been carried out from other monocot families like *Liliaceae, Amaryllidaceae, Alliaceae* and *Araceae*. Except for rice lectin, all Gramineae lectins have similar molecular structure, they are dimers composed of MW 18 kDa subunits, and recognize GlcNAc, oligomers of GlcNAc and GalNAc. Amarylidaceae lectins are dimers or tetramers composed of MW 13 kDa subunits, specific for mannose and mannose oligomers, they are mixtures of complex isolectins, and do not agglutinate human erythrocytes [8].

Lectin–carbohydrate interactions are crucial for the molecular events underlying the immune response. Lectins are therefore used as polyclonal reagents to investigate the molecular basis of lymphocyte activation and proliferation to identify and fractionate cells of the immune system. The first mitogenic agent described was Phytohemagglutinin (PHA), is a lectin isolated from *Phaseolus vulgaris* (red kidney bean) [9]. The discovery of lectin-mediated mitogenesis led to the detection of many other mitogenic lectins, most notably Concanavalin A [10], WGA [11], and Pokeweed mitogen (PWM) [12]. Some examples of mitogenic lectins include underground tubers of *Sauromatum guttatum* and *Gonatanthus pumilus* [13]. Lectins are also useful in detection of the alterations in the malignant cells hence used in cancer research and therapy [14]. In the present study we report for the first time the presence of lectins in monocot family *Aponogetonaceae* and characterized a novel lectin called *Aponogeton natans* tuber lectin (ANTL) and hemagglutinating, cytotoicity and mitogen proliferating potential of ANTL is evaluated.

## 2. Materials and methods

*Aponogetan natans* tubers are collected from the natural ponds of University if Hyderabad campus. All the reagents used were of highest available grade.

### 2.1. General analytical methods

The amount of protein was determined as described previously [15] using bovine serum albumin as standard. Total carbohydrate was estimated as described [16] using D-glucose as standard. Protein profile of the affinity chromatography fractions was monitored spectrophotometrically at 280nm. Hemagglutination assays were performed as described [17] with trypsinized human, rat and rabbit erythrocytes. SDS-PAGE was performed on a 1 mm thick vertical slab gel (10 cm x 10 cm).

### 2.2. Extraction and purification of ANTL

10 gm of fresh tubers were crushed in liquid nitrogen and proteins were extracted in 10 mM PBS buffer pH 7.2 at 4°C. The crude extract was centrifuged at 10,000 rpm at 4°C for 30 min, the supernatant was collected and filtered through whatman filter paper. The extract was saturated by gradient ammonium sulphate (0-80%) and incubated overnight by stirring at 4°C followed by centrifugation at 10,000 rpm at 4°C for 30 min, the collected pellet was dissolved in 10 mM PBS and dialyzed against PBS. Chitin flakes (Sigma) were crushed to fine powder by using mechanical blender. The affinity column was packed with fine powder. The final bed volume was maintained as 50 ml (14.5 × 75 mm) further, the column was washed by using 1000 ml of 0.1 M HCl followed by 1000 ml of 0.1 M NaOH and equilibrated with 10 mM PBS. The 15 ml of extracted sample was passed through the column until the absorbance is up to 0.02 at 280 nm and 3 ml fractions were collected, glacial acetic acid (0.2 M) was used for elution. The collected fractions were neutralized with Tris (1M), pooled further and dialyzed against PBS.

### 2.3. Molecular weight determination

The native molecular weight of lectin was determined by gel filtration chromatography using Sephadex G-200 column. The column was calibrated using proteins of known molecular weight i.e., Lysozyme (MW 14 kDa), ovalbumin (MW 45 kDa), Bovine serum albumin (MW 66 kDa), and Phosphorylase b (MW 90 kDa) and fractions (2.0 mL) were collected from gel filtration chromatography for further analysis.

### 2.4. Periodic acid-Schiff’s staining

Glycoprotein nature of the ANTL was determined by using periodic acid-Schiff’s staining as described previously [18]. Schiff’s reagent was prepared by adding 1gm of basic Fuschin to 200 ml of H_2_O by boiling at 70°C for few minutes followed by cooling and filtration. 2 gm of potassium metabisulphite and 5 mL of HCl was added to the schiff’s reagent and incubated overnight. 0.25 to 0.5 gm of activated charcoal is added to decolorize the solution completely and filtered. The staining solution was stored in stoppered brown bottle until further use. Purified HTNL protein was run on the SDS-PAGE and gel was incubated with 3% acetic acid and 1% periodic for one hour. After several washes with distilled water gel was stained in Schiff’s reagent for 30 min in dark. The gel was then destained in 10% acetic acid and incubated in 3% acetic acid for storage.

### 2.5. Stability studies

To determine the optimum temperature for ATNL activity, the ANTL protein (5 mg/mL) was incubated at different temperatures of 4 °C, 30 °C, 40 °C, 50 °C, 60 °C, 70 °C, 80 °C and 90 °C for 10 minutes. Hemagglutination activity of the protein samples was tested by using various erythrocytes. To determine the optimum pH required for the ANTL activity, the protein (5 mg/mL) was dissolved and incubated in buffers consisting different pH (20 mM sodium acetate/sodium phosphate, Tris-HCl) i.e., pH 4.0, 5.0, 6.0, 7.0, 8.0, 9.0 and 10.0 for a period of 30 min and the hemagglutination activity was analyzed. Haemaglutination activity was further analyzed with different concentrations of EDTA (10 mM – 30 mM).

### 2.6. Cytotoxic studies

SUP T1 cell lines (Non-Hodgkins T-Cell lymphoma) were obtained from Mc Kessan clinical and biological services, USA. U266 cell lines (Myeloma cell line) were obtained from national center for cell sciences, Pune. Both SUP T1, U266 cell lines were for the cytotoxicity studies.

Cultures in 96 well microtiter plates (200 µl) were incubated with various concentrations of ANTL (0-20 µg/ well) and centrifuged at 1000 rpm in room temperature.100 µl of the supernatant is discarded from the each well and 20 µl of 5 mg/ml MTT prepared in PBS was added to the wells. The plate was incubated at 37 °C for 4 hrs and 100 µl of SDS (0.01 N HCl in 10% SDS) was added to the each well. To solubulize the formazan compound, plates were incubated overnight. the absorbance of the each well was measured at 570 and 630 nm respectively.

### 2.7. Lymphocyte proliferation assay

Peripheral Blood Lymphocytes (PBL) were collected as described previously [19]. Heparinised tubes were used to collect venous blood (8U/ml) and diluted it to 1:1 with saline. 8ml of the diluted blood sample was transferred to 3 ml Histopaque and centrifuged for 20 min at 500 rcf. The cells at the interface were collected and suspended in the complete medium. The media used has the following composition: RPMI-1640 supplemented with 5% fetal calf serum and 50 mg/Liter Gentamycin, ^3^H-Thymidine comprising specific activity 5mCi/m mole. Scintillation cocktail consisting 4 gm each of PPO and POPOP in liter of Toluene has been used. 2 × 105 PBL and splenic lymphocytes in triplicate were cultured in 96 well flat-bottomed microwell plates with various concentrations of ANTL (0-20 µg), DNA, oligodeoxynucleotides with 200 µl of complete medium. Cultures were kept in a humidified incubator at 37 °C with 5% CO_2_. These cultures were treated with ^3^H-Thymidine (1µCi (for 24 hours and harvested on fiber filter using skatron automatic cell harvester. These filters were transferred to scintillation cocktail containing toluene. The radioactivity of the samples was measured using Beckman Scintillation Counter and the values were expressed as cpm/10^6^ cells.

### 2.8. Interaction of ANTL with normal murine, human Peripheral blood Lymphocytes and multiple myeloma cells

Flourescein isothiocyanate (FITC) labeled ANTL was prepared as described [20]. About 1% of the ANTL lectin in 0.025 M Na_2_CO_3_ and 0.025 M NaHCO_3_ was dialyzed against fluorescein isothiocyanate (0.1 mg/ml) in the same buffer. After 24 hr, the conjugated ANTL was dialyzed against PBS pH (7.3) until fluorescence was no longer detected in the dialysate. The conjugate was then ready for use. Human cancer cells (U266 & SUP T1 cells) and human peripheral blood lymphocytes and normal murine cells were incubated with FITC conjugated ANTL at room temperature for 15 min. The cells were then washed with PBS and finally suspended in 0.1 ml of the same buffer. The florescence in cells was observed under confocal scanning electron microscope with two sets of lymphocytes.

## 3. Results and Discussions

### 3.1. Purification and Electrophoretic analysis of ANTL

In the present investigation a chitin-binding lectin was isolated from the tubers of *Aponogeton natans* using chitin affinity chromatography on chitin column. Preliminary experiments with crude extract for haemagglutination demonstrated ANTL haemagglutinates different blood groups with different specificities. Haemagglutination studies done with the crude extract were inhibited by only chitin oligosaccharides (Table 2). Chitin column was used to purify the ANTL as it is shown specific inhibition by oligomers of *N*-acetylglucoseamine. The ANTL was eluted from the chitin column with 4-fold purification in a single step with yield up to 70.6% (Fig. 1). The results of affinity purification are summarized in Table 1.

**Table 1:** Purification table of ANTL.

| Sample | Protein (mg) | Specific activity*<br>(U/mg) | Total activity<br>(titre x mg) | Yield (%) | Purification factor |
| --- | --- | --- | --- | --- | --- |
| Crude protein | 104 | 524590 | 54557359 | 100 | 1 |
| Purified protein | 18 | 2133429 | 38399834 | 70.6 | 4 |

\* Specific activity is calculated by using highest dilution of protein which agglutinates with 2% Rabbit erythrocytes.

**Table 2:**
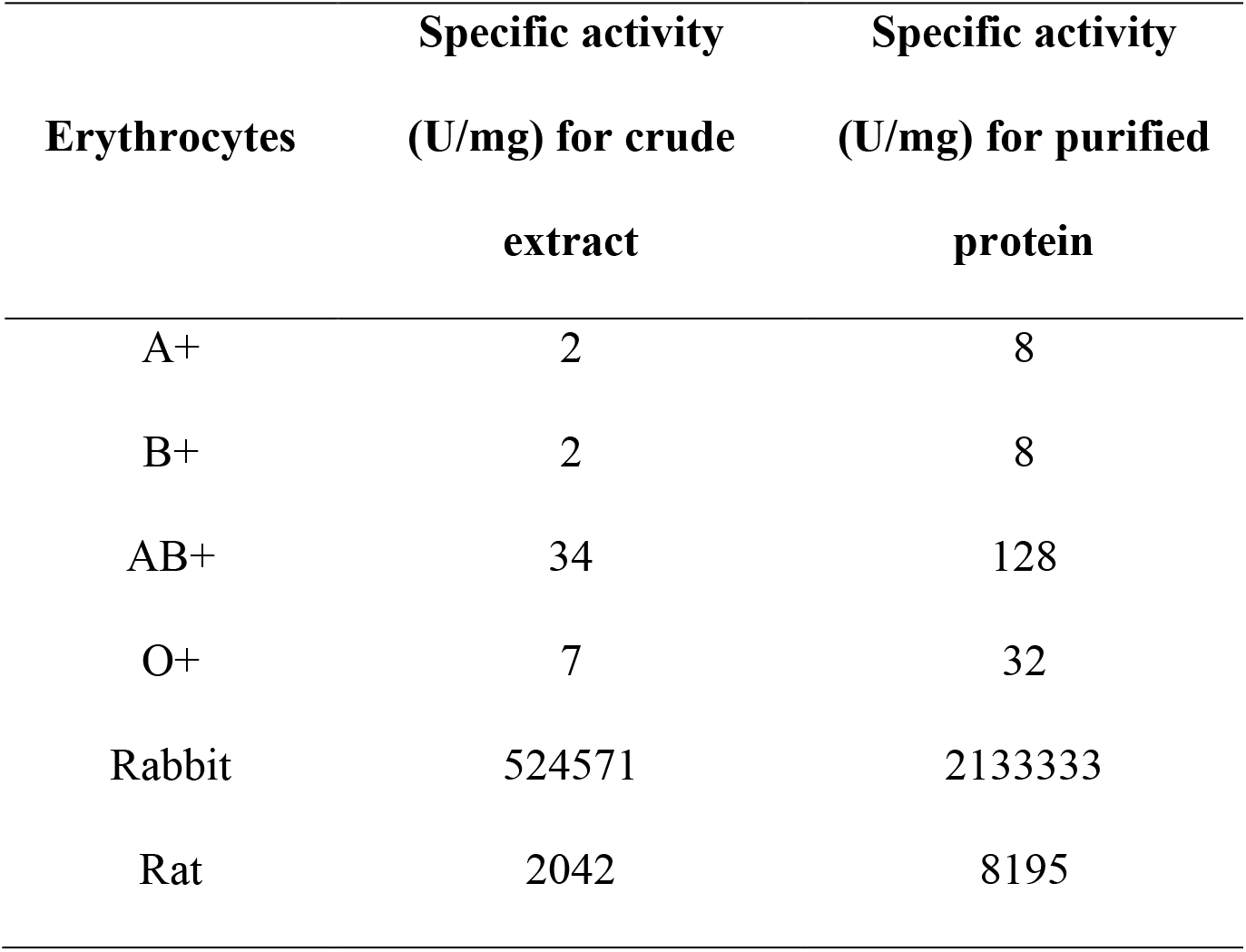
Erythrocyte specificity studies of purified lectins (ATNL)

**Figure 1:**
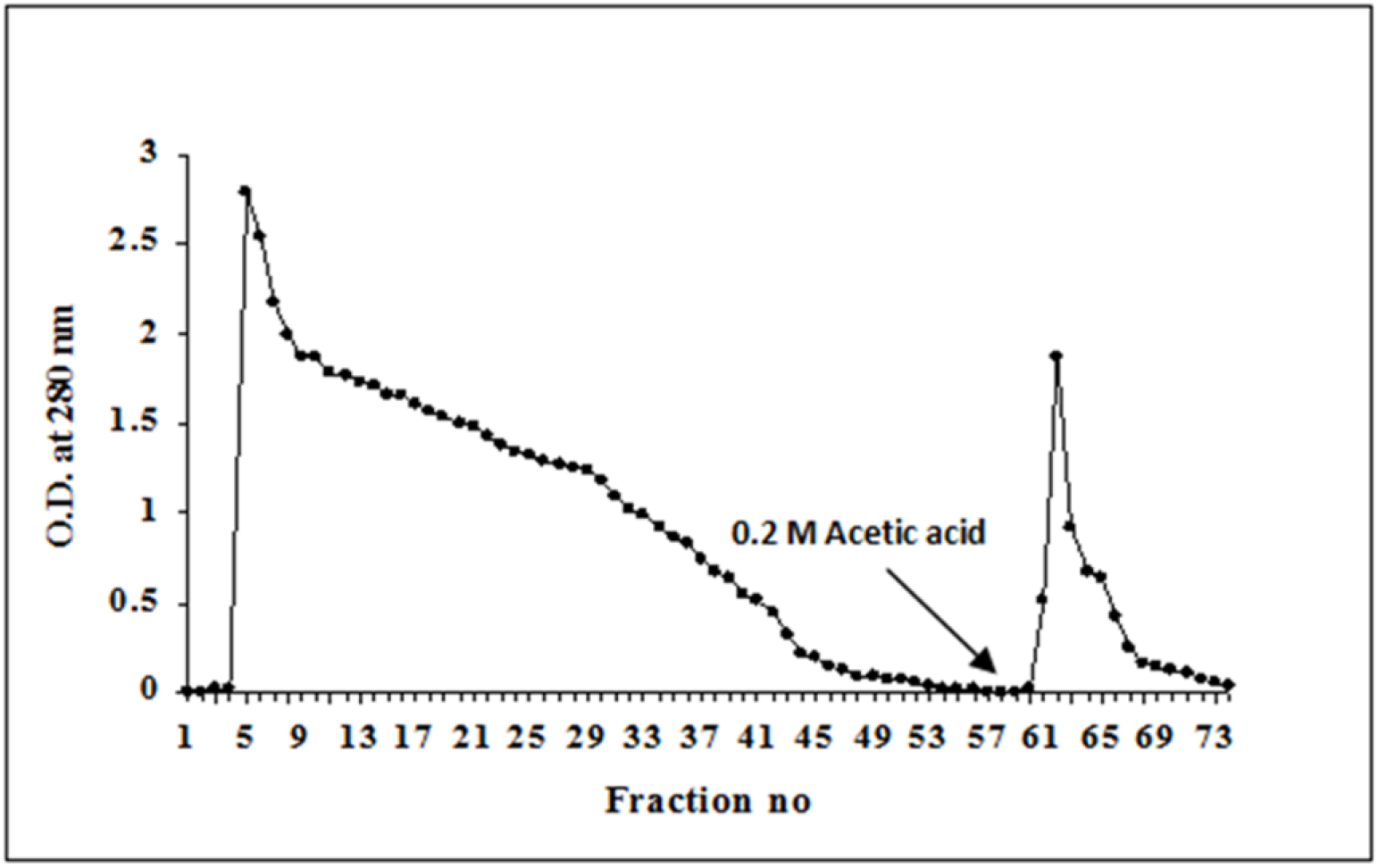
Affinity chromatography analysis of *Apanogeton natans* tuber lectin (ANTL) on chitin column. Crude tuber extract was passed through the chitin column. Individual fractions were eluted by using 0.2 M glacial acetic acid and collected. OD of each fraction was recorded at 280 nm, Peak in the graph showing the purified ANTL.

A reasonably high amount of lectin activity was recovered in the purified lectin. Although ANTL constituted only a small proportion of the total weight of tubers, interestingly they represented a considerable proportion of the tuber protein suggesting that such high lectin content may fulfill some physiological role in the plant survival [3]. The amount of total protein present was 2.7 mg/ml in crude extract and 0.33 mg/ml in chitin binding fractions. The lectin amount is 17.4% of the total protein in the tubers. The ANTL was resolved as a single band in the native PAGE which confirms the homogenity of the eluted lectin (Fig. 2A).

**Figure 2:**
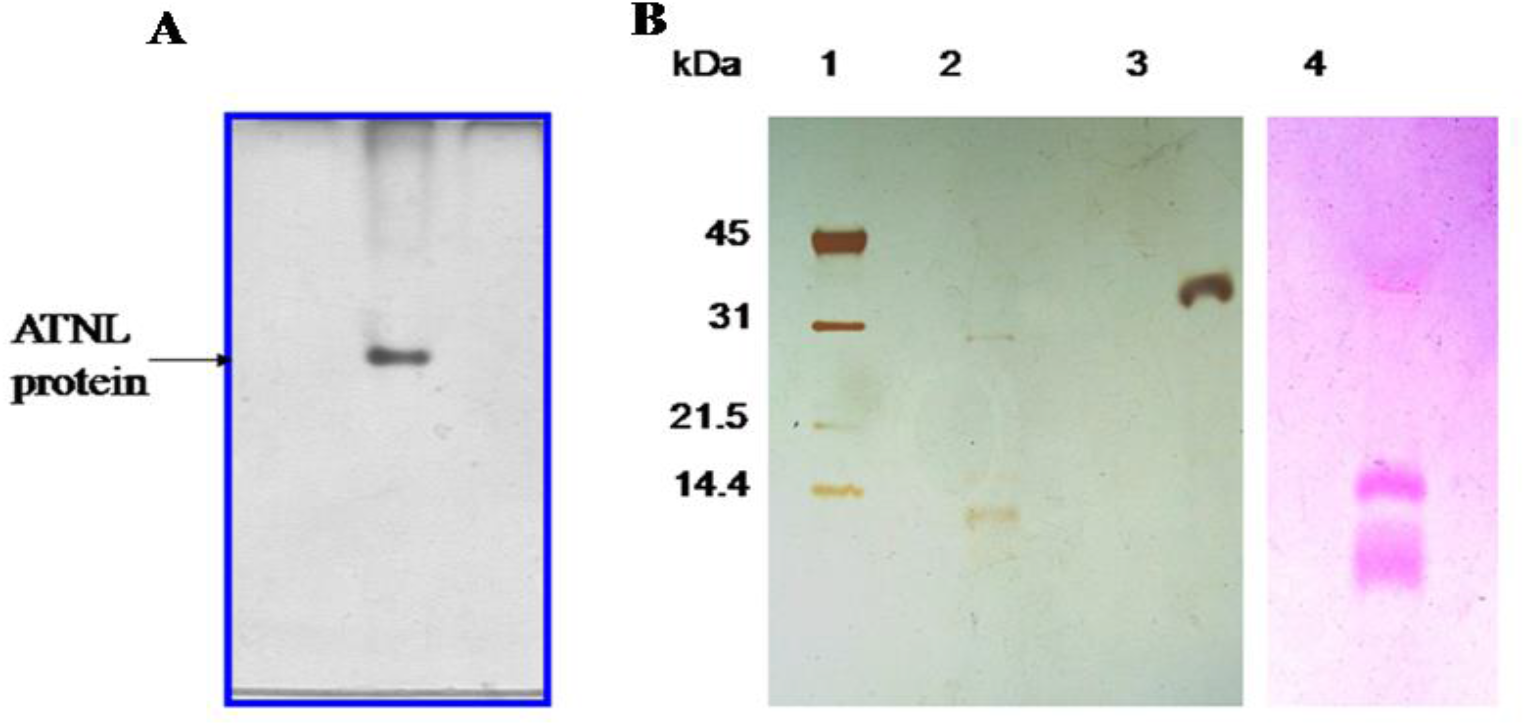
Native PAGE, SDS-PAGE and periodic acid staining (PAS) analysis of ANTL protein. **A)** Native PAGE analysis of ANTL protein. Collected fractions from affinity chromatography were run on the native PAGE gel. The ANTL was resolved into a single band in the native PAGE confirming the homogenity of the eluted glycoprotein. **B)** SDS-PAGE and periodic acid staining (PAS) of ANTL protein. Purified ANTL was separated by 12.5% SDS-PAGE. For PAS staining, the gel was stained in Schiff’s reagent for 30 min in dark followed by destaining in 10% acetic acid.

ANTL was resolved as a single band in SDS PAGE corresponding to the molecular weight of 33 KDa. Periodic acid staining confirmed the glycoprotein nature of the lectin in the gel. The carbohydrate amounts to 8.2% of the total protein (Fig. 2B). The apparent molecular weight of the pure protein is found to be 66 KDa as determined by the gel filtration chromatography.

ANTL was eluted as a single symmetrical peak with respect to the elution volume of standard bovine serum albumin suggesting it is a homodimer (Fig. 3).

**Figure 3:**
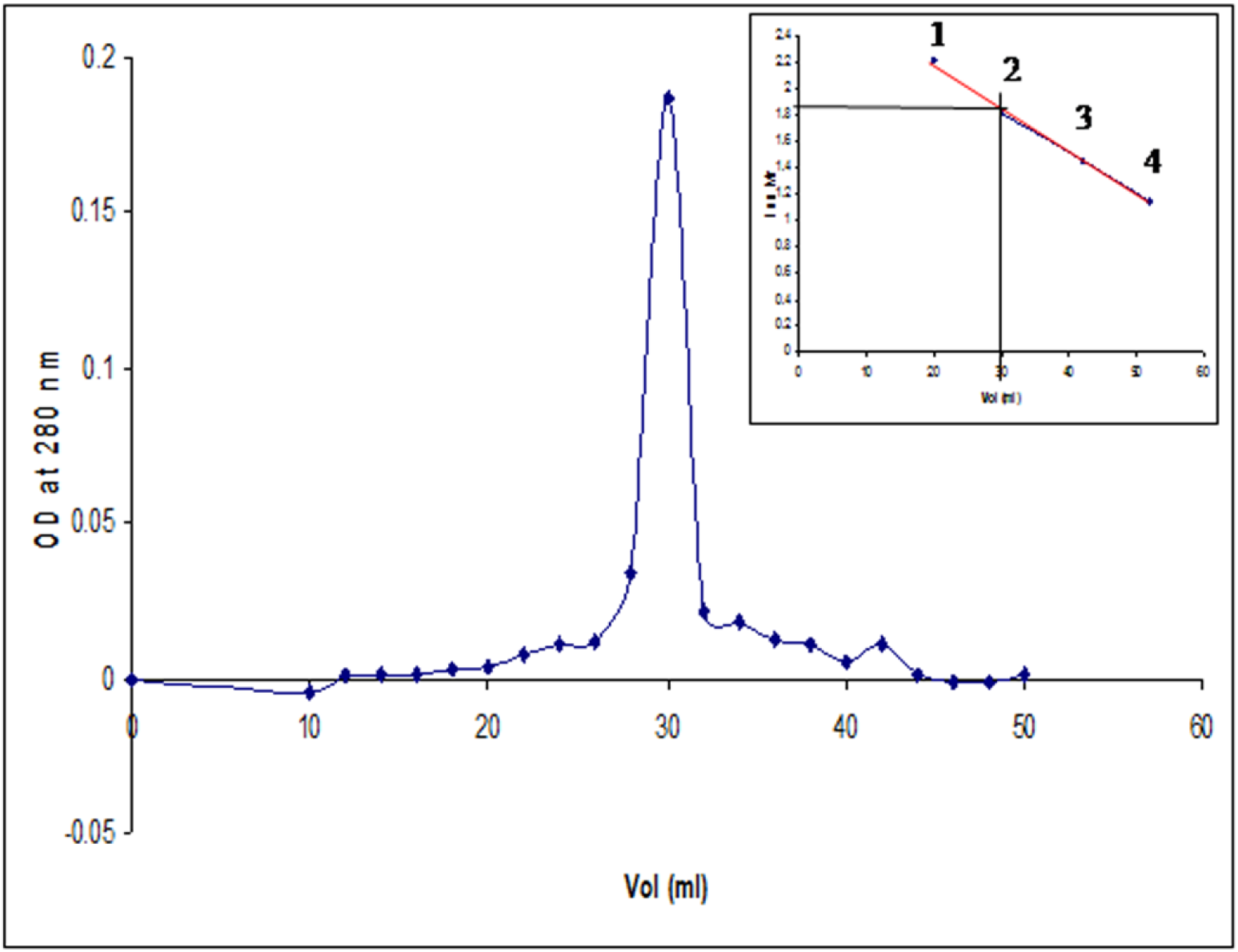
Molecular weight determination of ANTL protein by gel filtration chromatography (G-200). The native molecular mass of the ANTL was determined by using Sephadex G-200 column. ATNL molecular weight was calculated by calibration of the column with the known proteins i.e., Phosphorylase b (MW 90 kDa), BSA (66 kDa), ovalbumin (45 kDa) and Lysozyme (14 kDa).

### 3.2. Agglutination activity, sugar inhibition studies and stability analysis of ANTL

The ANTL lectin agglutinated RBCs from human, rabbit and rat. It has shown different specificities to different human blood groups i.e., A+, B+, AB+, O+, rabbit and rat. The ANTL has shown high specific activity on rabbit RBCs i.e., 2133333 U/mg which is highest till date reported in monocot lectins (Table 2).

The susceptibility of rabbit RBCs to higher agglutination suggests the presence of large number of available lectin receptors on rabbit RBC cells. Trypsinization has not altered the agglutination assay results suggesting that there is no unmasking on receptors upon trypsin treatment.

Lectin activity is assessed by checking its inhibitory potency by various sugars for haemagglutination activity. Glycoproteins and simple sugars did not inhibit the activity of lectin indicating the specificity of the lectin towards the *N*-acetylglucosamine oligomers. The agglutination activity of the lectin is only inhibited by *N*-acetylglucosamine in the following order: pentasaccharide > tetrasaccharide > trisaccharide > hexasaccharide > disaccharide (Table 3). The final concentration of the erythrocytes used in the study is 2% and the lectin is 2 haemagglutination Units (HU).

**Table 3:**
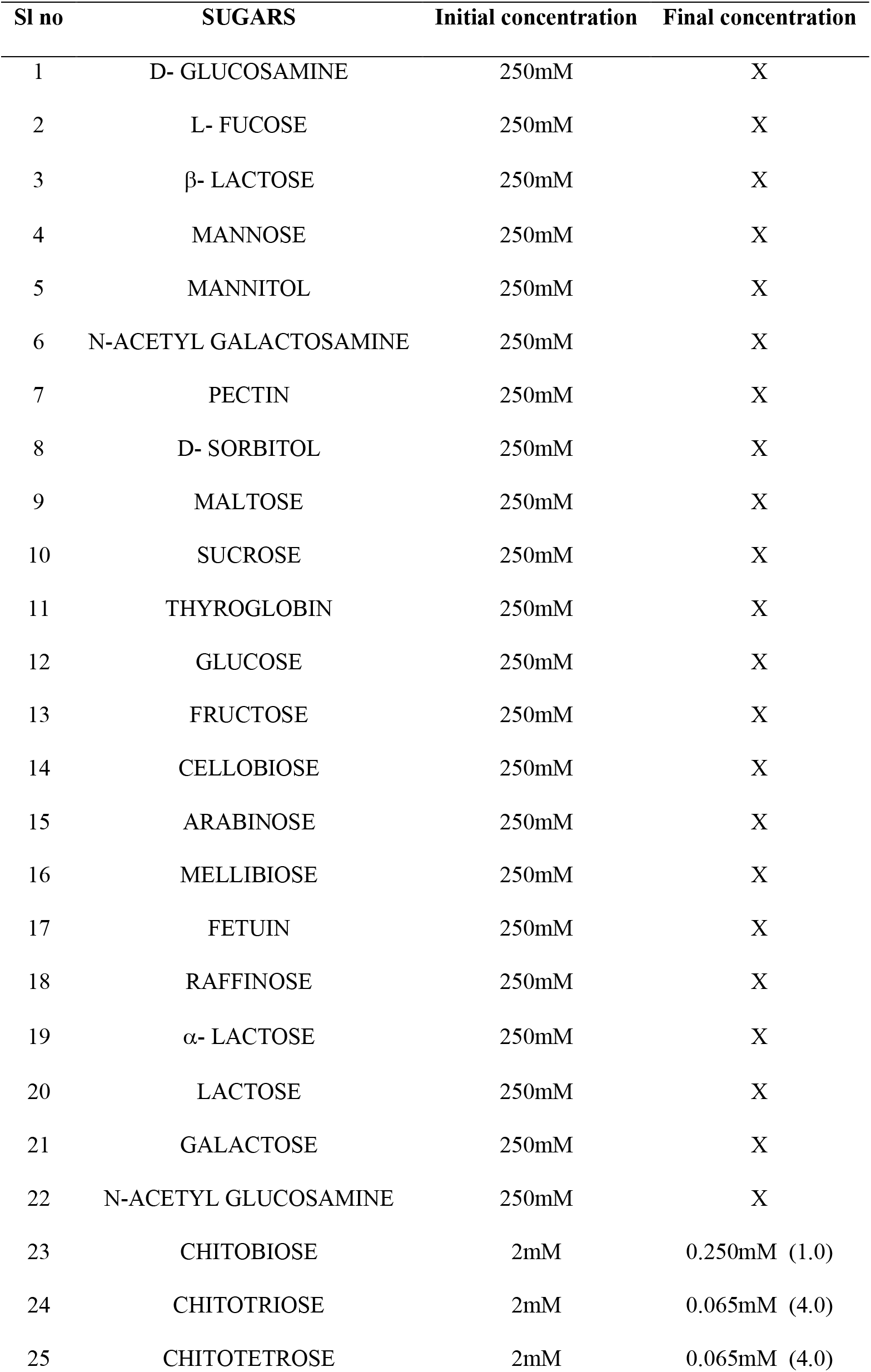
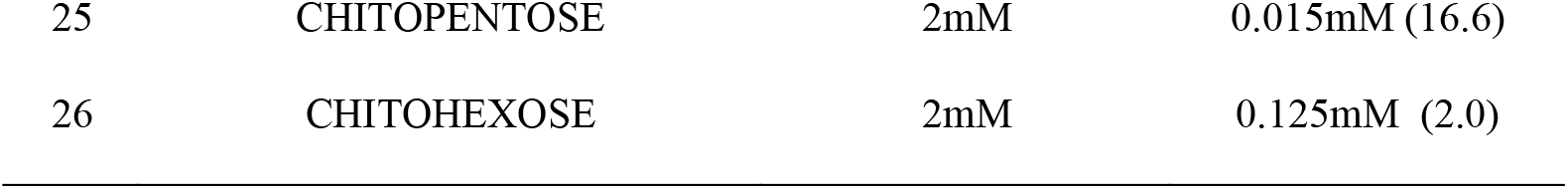
Sugar inhibition assay with various sugars and oligosaccharides.

Heat denaturation study of ANTL revealed that the protein is stable up to 60 °C and loses its activity at 90 °C upon treatment for 10 min. The ANTL has displayed maximum hemagglutination activity at pH 6 (Table 4). Low low carbohydrate content and the lack of disulphide bridges may be contributed to the low thermal stability of the HTNL. The lectin activity was affected by denaturing agents such as Urea (2M), Iodic acid (2M), Guanidium hydrochloride (2M), Lithium chloride (2M) and Potassium ferricyanide (2M), the agglutination activity was decreased with the increase in the concentration of the denaturing agents (data not shown). This decrease in lectin activity may be due to the disruption of hydrogen bonds and hydrophobic interactions of the ANTL by the denaturing agents. Hemagglutination activity of ANTL is not disturbed by EDTA (up to 30 mM) suggesting ANTL does not require metal ions for its activity. Similar pattern was observed in the case of *Amaryllidaceae* family members and other monocot lectins with respect to thermal stability, non-requirement of divalent metal ions and denaturing agents [21].

**Table 4:** Temperature optima and pH studies with rabbit erythrocytes.

| Temperature<br>(°C) | Specific activity<br>(U/mg) | pH | Specific activity<br>(U/mg) |
| --- | --- | --- | --- |
| 30 | 2133333 | 4 | 522448 |
| 40 | 2133234 | 5 | 2133261 |
| 50 | 522448 | 6 | 2133333 |
| 60 | 8195 | 7 | 522448 |
| 70 | 32 | 8 | 2051 |
| 80 | 16 | 9 | 2051 |
| 90 | 8 | 10 | 2051 |
| 100 | 0 | 11 | 2051 |

### 3.3. Cytotoxic and Mitogenic proliferation studies on ANTL

The cytotoxic effect of the ANTL was studied using mammalian cell lines SUP T1 and U266. After 72 hrs of incubation SUP T1 cell lines with ANTL, they have not shown any significant decrease in the proliferation. Whereas, U266 cell lines treated with ANTL have displayed ∼50 % inhibition in the proliferation with respect to the controls and a gradual decrease in the proliferation of U266 cell lines has been observed with increasing concentration of the ANTL protein (Table 5).

**Table 5:**
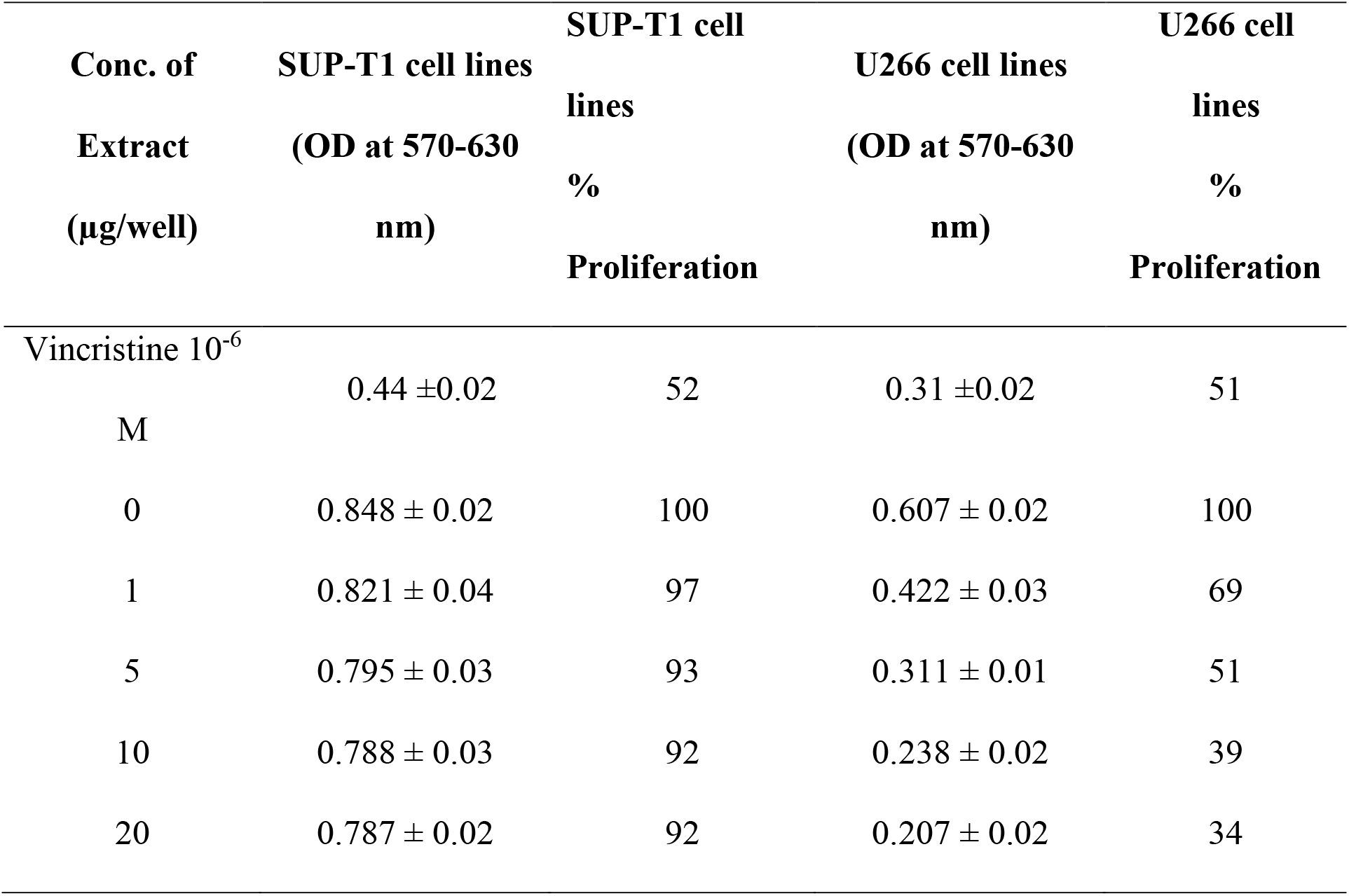
Proliferative response of SUP-T1 and U266 cell lines treated with ANTL.

The variation of proliferation inhibition of ANTL on mammalian cell lines SUP T1 and U266 may be due to the different glycoconjugates present on the surface of these cell lines, thus leading to the different downstream signaling events of lectins. As every lectin display specificity to its interaction with particular sugar, there is a need to check a wide range of lectins against various cancer cell-lines. The exact molecular mechanism(s) behind the plant lectins involvement in the anti-proliferative effect of human cell lines is not clear at present. Several hypotheses suggests that this may be due to the ability of lectins to modulate the growth, proliferation, differentiation and apoptosis of mammalian cells both in vivo and in vitro conditions. Additional studies are required to understand mechanism of antiproliferative activity plant lectins on mammalian cell lines.

As it is known that *Euphorbiaceae, Leguminaceae, and Gramineae* lectins acts as a potent mitogen on normal splenic lymphocytes, the mitogenic activity of ANTL was analyzed on murine and human splenic lymphocytes. ANTL has exhibited mitogenic response towards both murine and human lymphocytes as evidenced by lymph proliferation in the cultures upon treatment with ANTL. The mitogenic stimulation of ANTL on murine and human lymphocytes was significantly higher than that of a well-known standard plant mitogen Con A. The optimum proliferation dose of ANTL was 1 µg/ml for murine and 5 µg/ml for human lymphocytes (Table 6).

**Table 6:**
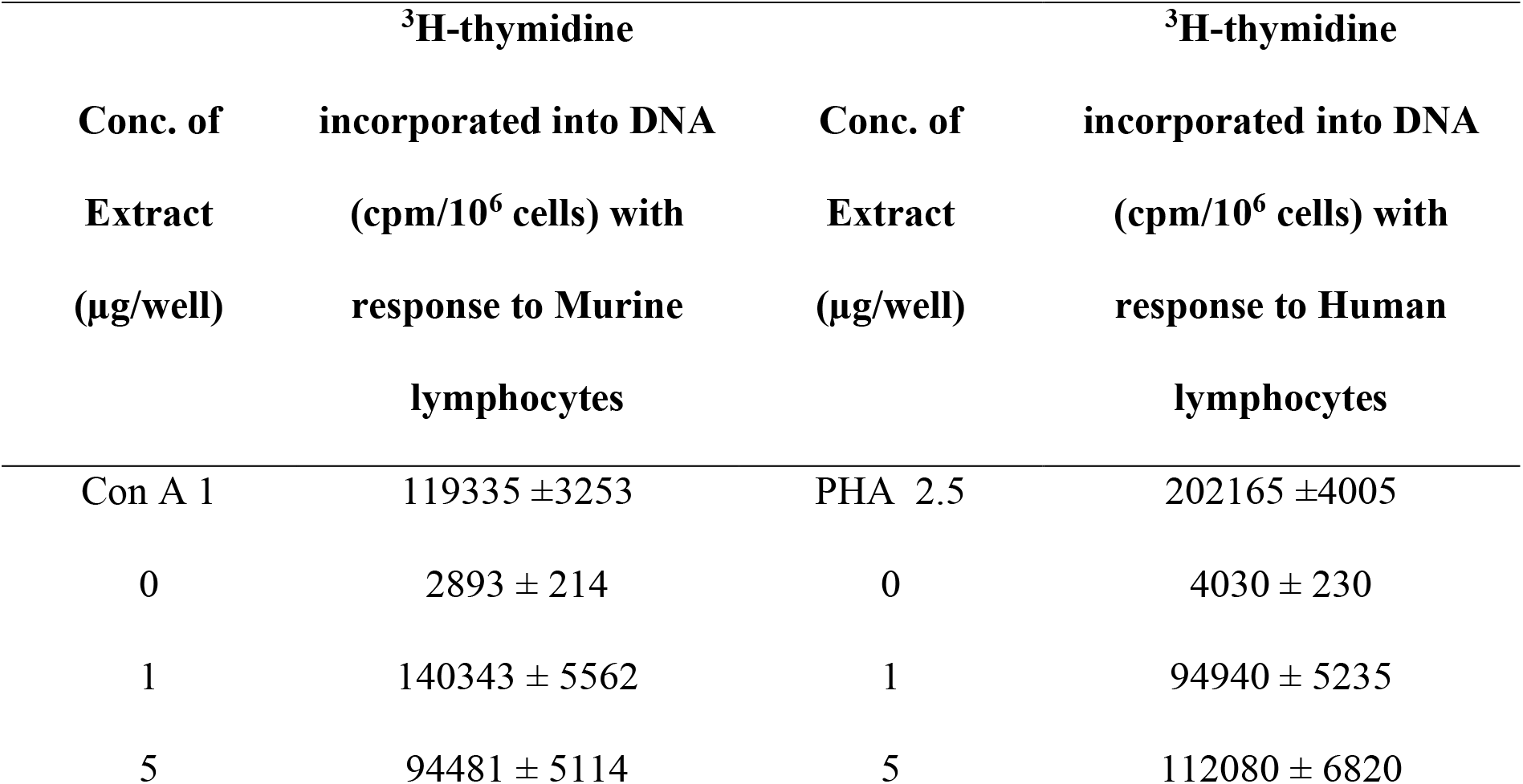
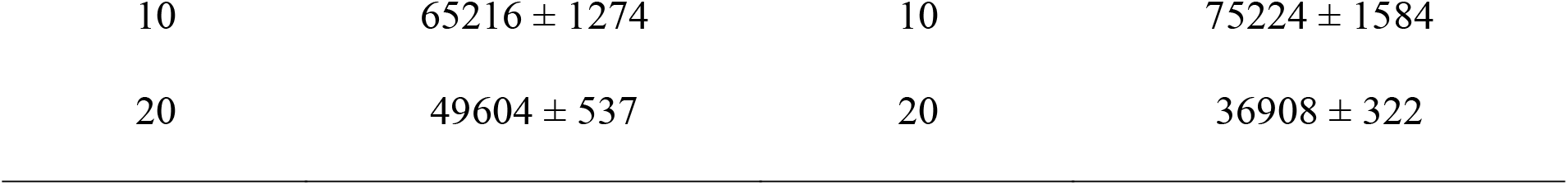
Proliferative response of ANTL on Murine and Human lymphocytes.

The determination of optimum concentration of the lectin is essential for mitogenic studies as higher lectin concentrations inhibit mitogenesis as evidenced by our study (Table 6) Mitogenic assays are mostly carried on lymphocytes as they are usual target cells for proliferation studies. The study of lectin–lymphocyte interactions lead to the understanding of mechanism of lymphocyte activation and their regulation, thus contributing to the better understanding of cell growth and development. The mitogenic response of ANTL protein on murine and human lymphocytes was inhibited in the presence of chitohexose in a concentration dependent manner. The inhibition of mitogenicity and hemagglutination of the HTNL in the presence of *N*-acetylglucosamine suggest that HTNL is involved in these processes by binding to the cell membrane receptors, which is recognized and inhibited by chitohexose like structures on the cells. Increase in the concentration of chitohexose reduced binding of lectin to the human and murine lymphocytes due to the decreased in the available sugar binding sites on lectin. FITC-conjugated lectin studies revealed that the lectin is able to bind with SUP-T1 T cell-lymphoma cells, U266 myolema cell lines, murine lymphocytes and human peripheral blood lymphocytes. However, the exact mechanism with which it is able to induce responses only in in some cell lines is not clearly understood (Fig. 4). Overall, mitogenic lectins can be used to study the relationship between chromosomal abnormality and human diseases, which eventually help in the diagnosis and treatment of diseases.

**Figure 4:**
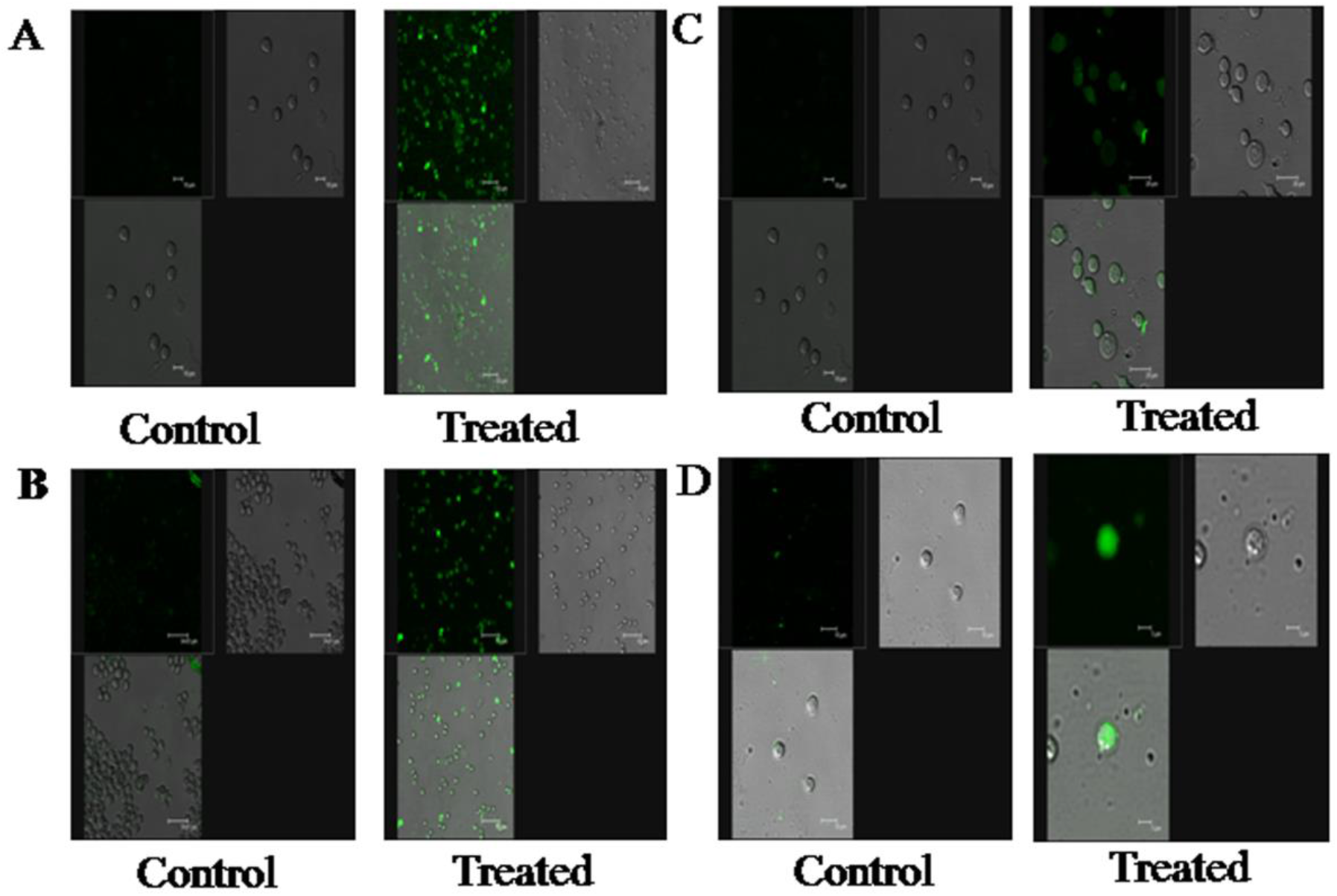
Interactions of FITC-Conjugated ANTL with A) SUP-T1 T Cell-lymphoma B) U266 myolema cell lines C) Murine lymphocytes and D) Human peripheral blood lymphocytes.

## 4. Conclusions

The current investigation revealed the presence of a chitin-binding lectin from the tubers of a monocot species *Aponogetonnatans* (ANTL). For the first time we have reported the presence of lectins from the family *Aponogetonaceae*. Lectin activity was inhibited by the chitin *N*-acetylglucosamine. ATNL was isolated and purified by Affinity chromatography on chitin column followed by gel filtration chromatography. ANTL is a dimeric glycoprotein with a molecular weight of ∼66 kDa as revealed by Native PAGE and gel filtration chromatography analysis and ATNL exhibited its glycoprotein nature on periodic acid–Schiff’s reagent gel. Lectin consist 8.2% carbohydrate content. Unlike other lectins, ANTL protein was thermo stable up to 50°C with broad pH optima (pH 4–10). Functional analysis of ATNL revealed that it hemagglutinates rabbit erythrocytes as well as human blood cells of groups A, B and O with different specificities. Mitogenic activity study revealed that ATNL induces the proliferation on human lymphocytes and murine cells at the concentration as low as 1 µg/ml. Upon treating human U266 cells with HTNL, they have exhibited 50% decrease in the proliferation. Hemagglutination and mitogenic proliferation activity of ANTL suggests that it is a Chitin-binding lectin with diverse functions. Overall, the present study advances our knowledge of monocot lectins and insights into their functions.

## Conflict of interest

Authors declare that they have no conflict of interest.

## Acknowledgements

We acknowledge the financial assistance from CSIR, New Delhi, India. The authors thankful to Department of Plant Sciences, School of Life sciences, University of Hyderabad for facilities.

## Notes

### Competing Interest Statement

The authors have declared no competing interest.

